# Machine Learning Classification of Alzheimer’s Disease Pathology Reveals Diffuse Amyloid as a Major Predictor of Cognitive Impairment in Human Hippocampal Subregions

**DOI:** 10.1101/2023.05.31.543117

**Authors:** T.L. Stephen, L. Korobkova, B. Breningstall, K. Nguyen, S. Mehta, M. Pachicano, K.T. Jones, D. Hawes, R.P. Cabeen, M.S. Bienkowski

## Abstract

Analyzing Alzheimer’s disease (AD) pathology within anatomical subregions is a significant challenge, often carried out by pathologists using a standardized, semi-quantitative approach. To augment traditional methods, a high-throughput, high-resolution pipeline was created to classify the distribution of AD pathology within hippocampal subregions. USC ADRC post-mortem tissue sections from 51 patients were stained with 4G8 for amyloid, Gallyas for neurofibrillary tangles (NFTs) and Iba1 for microglia. Machine learning (ML) techniques were utilized to identify and classify amyloid pathology (dense, diffuse and APP (amyloid precursor protein)), NFTs, neuritic plaques and microglia. These classifications were overlaid within manually segmented regions (aligned with the Allen Human Brain Atlas) to create detailed pathology maps. Cases were separated into low, intermediate, or high AD stages. Further data extraction enabled quantification of plaque size and pathology density alongside ApoE genotype, sex, and cognitive status.

Our findings revealed that the increase in pathology burden across AD stages was driven mainly by diffuse amyloid. The pre and para-subiculum had the highest levels of diffuse amyloid while NFTs were highest in the A36 region in high AD cases. Moreover, different pathology types had distinct trajectories across disease stages. In a subset of AD cases, microglia were elevated in intermediate and high compared to low AD. Microglia also correlated with amyloid pathology in the Dentate Gyrus. The size of dense plaques, which may represent microglial function, was lower in ApoE4 carriers. In addition, individuals with memory impairment had higher levels of both dense and diffuse amyloid.

Taken together, our findings integrating ML classification approaches with anatomical segmentation maps provide new insights on the complexity of disease pathology in AD progression. Specifically, we identified diffuse amyloid pathology as being a major driver of AD in our cohort, regions of interest and microglial responses that might advance AD diagnosis and treatment.

## Introduction

The hippocampus is one of the first brain regions affected by AD and degeneration of hippocampal neurons is believed to underlie AD cognitive symptoms^1^. Recent work has highlighted AD as a heterogeneous disorder with numerous phenotypes and genotypes that ultimately progress along a continuum toward a similar clinical and pathological endpoint^2^. Generally, NFTs have a stereotypical progression pattern in AD: first appearing in the entorhinal cortex (Braak Stage I), then spreading into CA1 of the hippocampus (Braak Stages II-IV), before distributing throughout the rest of the hippocampus and cortical and subcortical brain areas (Braak Stages V-VI)^3^. Amyloid pathology has its own distinct, but highly variable progression pattern (Thal Staging), which is difficult to correlate with disease state and cognitive deficits^4,5^. Additionally, microglia undergo varying regional dependent functional changes, which can exacerbate AD cognitive decline, but the full clinical relevance remains unclear^6,7^. Overall, the varying clinical expression and pathophysiology has led to multiple AD subtypes being broadly described^2,8–17^, but how differences in AD heterogeneity emerge at subregional and cellular resolution is not known. Indeed, several different morphological types of AD pathology with distinct molecular compositions and glial interactions have been described, but it is not evident which pathological subtypes are most associated with clinical disease symptomatology.

Due to their physical and structural differences, artificial neural networks can be trained to identify and classify distinct pathological aggregates^18,19^. However, most ML approaches do not consider the anatomical distribution of these aggregates, which is important to understanding their impact on brain structure and function in AD^20,21^. Ultimately, ML approaches that can both classify distinct types of pathology and assess their distribution within distinct anatomical subregions are needed to characterize AD heterogeneity. The main aim of our study was to further understand neuropathological heterogeneity at the cellular level in distinct hippocampal subregions and layers. To do this we created the first combined neural network classification, segmentation and quantification approach to generate anatomically accurate pathology maps and investigate the spatial distribution of morphologically distinct types of amyloid, NFTs and microglia. Subsequently, aligning this data with cognitive outcomes and delineating ApoE and sex differences could have critical relevance in the field of AD diagnostics.

## Materials and Methods

### Case Cohort Imaging and Pathology Categorization

Posterior coronal consecutive sections (at the level of the lateral geniculate thalamus) stained with 4G8 antibody (amyloid) and Gallyas silver stain (NFTs) from 51 de-identified patients were obtained from the USC ADRC (Table 1) and imaged using a slide scanning microscope (Olympus VS120, 20X magnification) to create digital whole slide images. We grouped cases into low, intermediate and high based on the National Institute on Aging–Alzheimer’s Association guidelines for the neuropathologic assessment of Alzheimer’s disease^22^.

**Table 1.** Demographics of Low, Intermediate and High AD Cases. Age of death and postmortem intervals are presented as Mean (SD). Sex (male, female), ApoE (e4 carriers, e4 non-carriers) and cognitive status (mild-low, impaired) are presented as %.

|  | **All (n = 51)** | **Low (n = 14)** | **Intermediate (n = 18)** | **High (n = 19)** |
| --- | --- | --- | --- | --- |
| Age, mean years (SD)* | 84.8 (9.8) | 88.1 (7.0) | 88.0 (10.0) | 79.5 (9.3) |
| Sex (Male/Female, %) | 45/55 | 43/57 | 61/39 | 58/42 |
| APOE (4+/4-, %) | 47/53 | 31/69 | 47/53 | 58/42 |
| PMI, mean hours (SD) | 11.7 (12.7) | 11.4 (14.4) | 11.5 (10.4) | 12.3 (13.9) |
| CDR Memory Score (Low/Impaired, %)** | 31/69 | 82/18 | 29/71 | 0/100 |

### Image Annotation and Segmentation

4G8 sections were downsampled with a factor of 16 with Imagemagick and manually annotated and segmented with reference to the closest matching Allen Human Brain Atlas. Anatomical segmentation maps of the hippocampus and medial temporal lobe were manually delineated based on hematoxylin counterstain in 4G8 stained tissue sections using Quantitative Imaging Toolkit (QIT) software^23^ mask drawing module. QIT is a software package of computational tools for the modeling, analysis, and visualization of scientific imaging data. Regions of interest (ROIs) included Brodmann Area 35 and 36 (A35 and A36), CA1-4, dentate gyrus (DG), medial entorhinal cortex (MEC), parasubiculum (PAS), prosubiculum (ProR), presubiculum (PrSr), and subiculum (Sub). For all cases, corresponding Gallyas stained sections were aligned to 4G8 stained sections using Advanced Normalization Tools (ANTs) Quick Registration algorithm. Every section was checked and manually corrected with a warping module in Adobe Photoshop (Version: 22.3.0 20210302.r.49 660cd2e x64).

### Machine Learning and Classification

The ML toolkit Ilastik (interactive learning and segmentation toolkit) was utilized to classify 4G8, Gallyas and Iba1 stained images. Ilastik uses a random forest classifier in the learning step to characterize each pixel’s neighborhood with a set of generic (non-linear) features. Ilastik provides feedback in real time, enabling refinement of the segmentation result, to fine-tune the classifier with an uncertainty measure guiding the user to ambiguous regions in the images. Once a classifier has been trained on a set of representative images, it can be used to automatically process a very large number of images. To perform a more robust analysis of our 4G8 signal, we classified and segmented 3 types of amyloid pathology: individual amyloid particles (4G8 labeled intracellular APP, extracellular dense plaques, and extracellular diffuse amyloid. Due to blood vessel staining in the Gallyas signal, Ilastik was used to create layers with blood vessels and NFTs respectively. The layer containing blood vessel pixels was then removed for further processing. Other artifacts, such as tissue tears, were manually erased in Photoshop to ensure high specificity for NFTs. By co-registering Gallyas and 4G8 stained images, we were also able to identify and quantify the distribution of neuritic plaques (defined as 4G8 amyloid and Gallyas NFT colocalized plaques). Region map segmentation results were visualized and corrected semi-automatically in QIT.

### Quantification and Statistical Analyses

Ilastik-processed image data extracted with QIT was co-registered to the corresponding segmentation map to determine pathology classified pixels within anatomically segmented subregions. We computed the pixel-wise overlap to detect the anatomical regions coincident with four types of pathology signal: Gallyas positive NFTs, dense amyloid, diffuse amyloid, and amyloid speckles (APP). Each mask was treated separately and two quantification procedures were performed. In the first approach, the total number of pixels for each pathology type in each annotated ROI was extracted and densities calculated based on region areas. In the second approach, we looked at clusters of pathology using an algorithm for connected components, which provided a complete sampling of the distribution of pathology cluster sizes for each anatomical region and pathology type. Statistical analyses were performed using RStudio (Version 1.3.959). ANOVA was run for sex, stage and ApoE interactions accordingly and Wilcox post-test was applied for non-parametric data.

### Cognitive Data

Clinical Dementia Rating (CDR) Dementia Staging Instrument Plus National Alzheimer’s Coordinating Center (NACC) frontotemporal lobar degeneration (FTLD) Behavior and Language Domains, using only the memory part, were classified as 0.0 (no impairment), 0.5 (questionable impairment), 1 (mild impairment), 2 (moderate impairment) and 3 (severe impairment). We combined 0, 0.5 and 1 as mild/no impairment and 2 and 3 as impaired.

## Results

### Pathology classification within medial temporal lobe sections

To analyze and extract robust pathology data from USC ADRC brain tissue sections, a ML based pipeline was established to classify and quantify distinct types of AD pathology within neuroanatomical subregions (Fig. 1a). To differentially characterize anatomical features of AD pathology, Ilastik was trained to classify different features of histological labeling. Extracellular amyloid deposits were labeled using 4G8 antibody staining, which also cross-reacts to label intracellular amyloid precursor protein APP^24^. Classifications included intracellular APP, dense plaques, and diffuse amyloid in 4G8 sections, NFTs in Gallyas sections, and neuritic plaques through colocalization of 4G8 and Gallyas-positive pixels in co-registered images (Fig 1b). Diffuse amyloid was defined as loose 4G8 staining with irregular, poorly defined margins (Fig 1b4) while dense amyloid plaques had clear outlines and generally had a strong, compact staining (Fig 1b5). Dense Aβ plaques are extracellular deposits composed of Aβ peptide, largely amyloid filaments, while in diffuse plaques, most of the protein is not aggregated as amyloid filaments. APP was defined as small intracellular deposits of 4G8 (Fig. 1b6) and NFTs are Gallyas positive intracellular tau deposits (Fig 1b10). Classified pixels of all AD pathology types were co-registered and re-colored to create new image maps of AD pathology distribution (Fig. 1c).

**Fig. 1.**
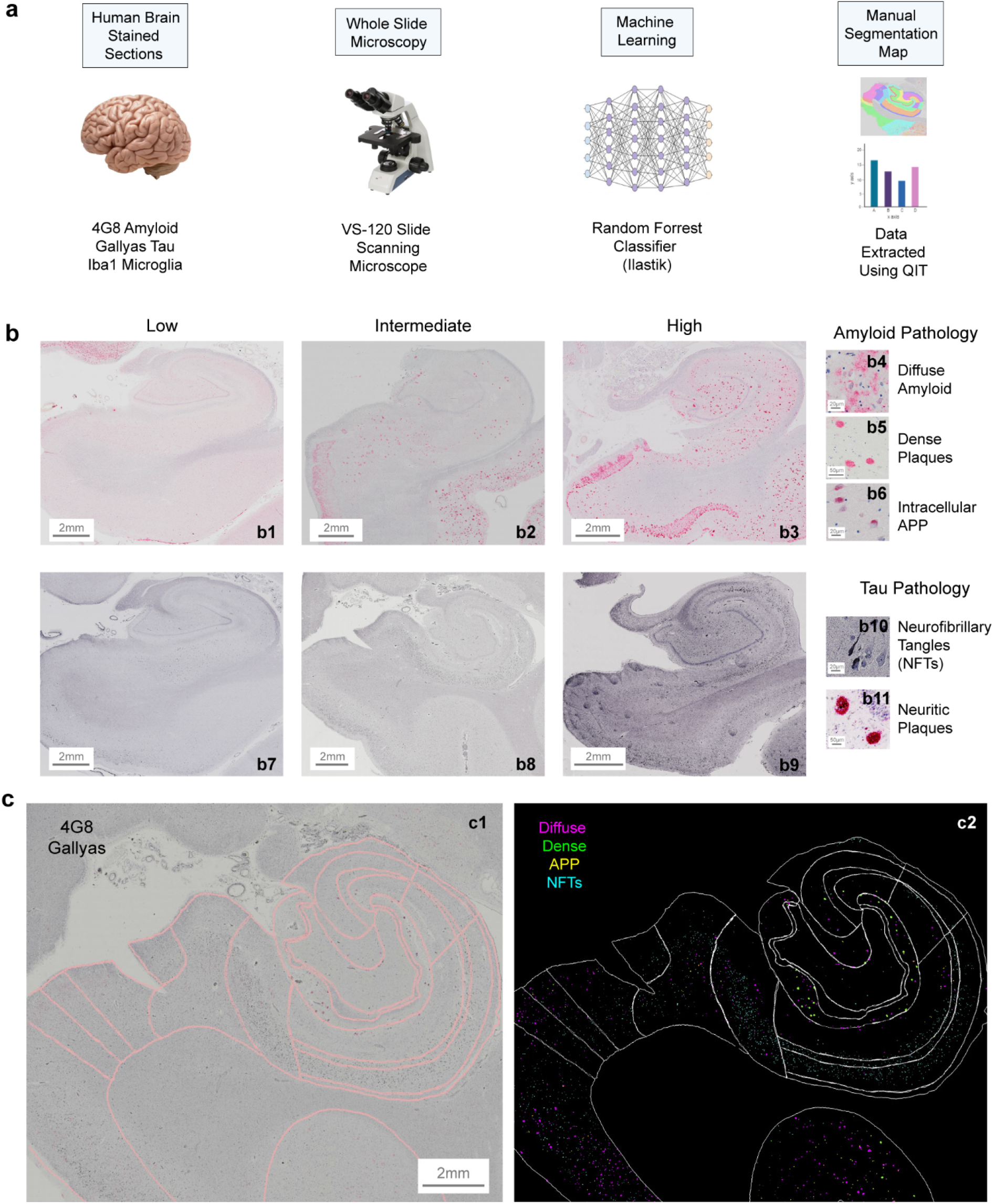
Hippocampal digital pathology analysis. Postmortem sections stained for amyloid (4G8), NFTs (Gallyas) and Iba1 microglia were imaged using a slide scanning microscope and analyzed using ML random forest classification, followed by segmentation and data extraction with QIT (a). Low, intermediate and high amyloid (b1-3) and NFTs examples (b7-b9) with example classification of amyloid plaques (diffuse, b4; dense, b5; intracellular APP, b6), NFTs (b10) and neuritic plaques (NFTs and amyloid positive, b11). Raw 4G8 amyloid signal (c1) with segmented pixels (c2).

In a subset of cases (n = 47), we then validated our approach to detect dense amyloid plaques in whole 4G8 sections using QIT software^23^ against the more established bioimage analysis software program QuPath and found these results were significantly correlated (Fig. 2a, *R* = 0.71, p < 0.0001). Based on the total amyloid pathology quantified within each AD patient, patients were grouped into low, intermediate, and high AD groups based on our sample dataset. Quantification of whole sections from 51 AD patients revealed that all three amyloid pathology types increased across disease stages with diffuse amyloid plaques increasing most rapidly within the hippocampus and medial temporal lobe (Fig. 2b). Overall diffuse amyloid was higher than dense and APP (Fig. 2b, type F2,147 = 14.76, p < 0.0001). In high cases diffuse staining was much higher than dense and APP (diffuse vs APP W = 68, p = 0.0023; diffuse vs dense W = 88, p = 0.019). All types of amyloid pathology increased with disease stage (Fig. 2c, stage F_2,299_ = 60.6, p < 0.0001) with a significant interaction between stage and pathology type (Fig. 2b-c F_2,902_ = 57.1, p < 0.0001). The number of dense amyloid plaques was significantly higher in high cases compared to low (Figure 2d, W = 162, p = 0.032) although we did not find any differences in plaque size (Supplementary Fig. 1c).

**Fig. 2.**
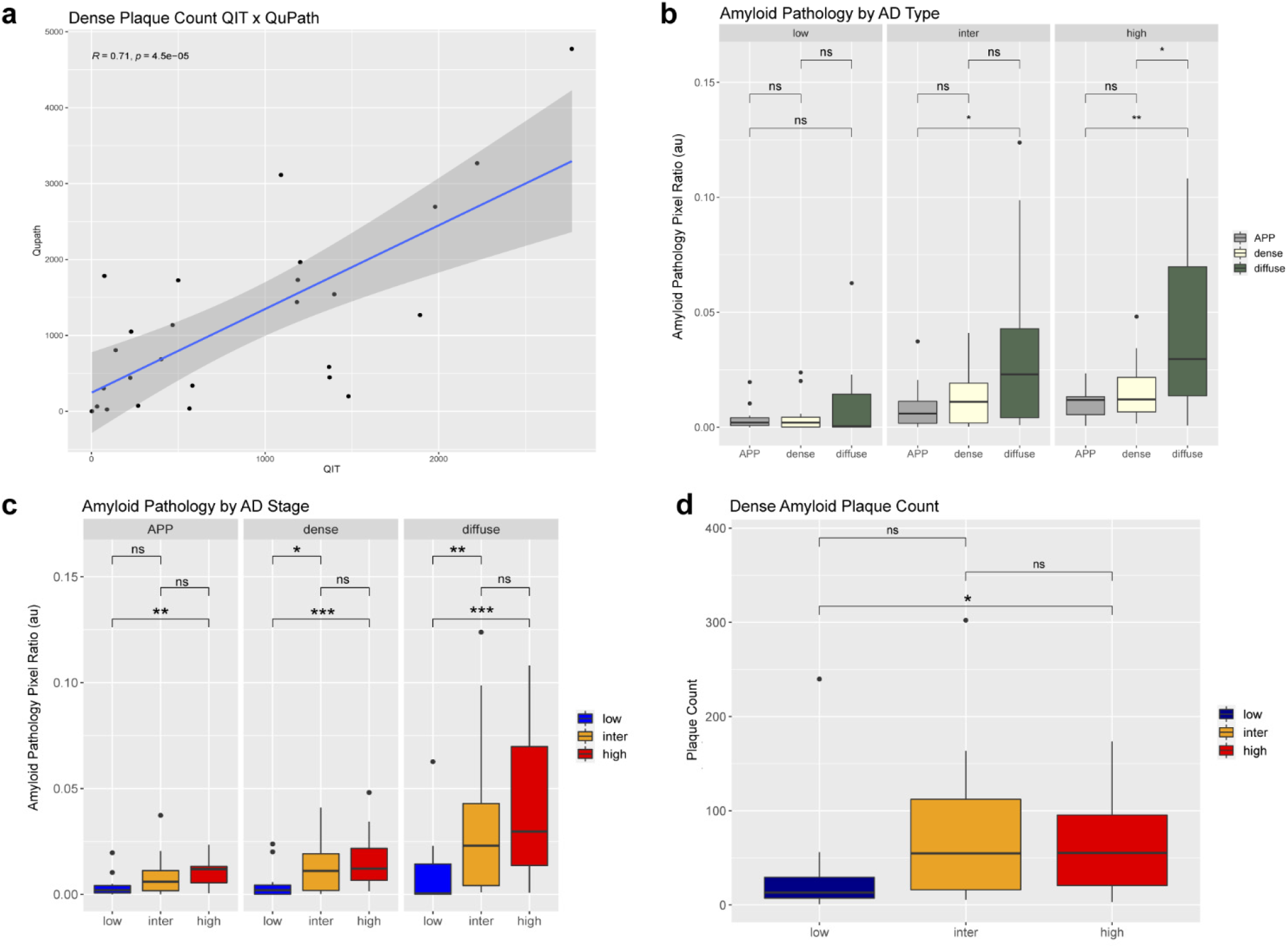
Data extracted with QIT and QuPath from 47 subjects (a). Amyloid pathology from QIT comparing dense, diffuse and APP (b). Amyloid pathology across low, intermediate and high cases (c). Dense plaque counts (d). Box plots show median (box center line), interquartile range (IQR, bounds of box), minimums and maximums within 1.5 times the IQR (whiskers), and outliers (points beyond the whiskers). *p* ≥ 0.05 was considered not significant (ns); \**p* < 0.05, \*\**p* < 0.01, \*\*\**p* < 0.001.

### Quantification of pathology types within hippocampal subregions

By registering our data with the Allen Human Brain Atlas, 4G8 sections were manually annotated and segmented using QIT into layered ROIs: Brodmann Area 35 and 36 (A35 and A36), CA1-4, dentate gyrus (DG), medial entorhinal cortex (MEC), parasubiculum (PAS), prosubiculum (ProR), presubiculum (PrSr), subiculum (Sub) and white matter (WM), which was included as a control region (Fig. 3a1). Total amyloid pathology pixels were subsequently aligned with these regions and normalized to region area to calculate pathology density (Fig. 3a2). In all hippocampal subregions, amyloid pathology increased from low to intermediate stage and most subregions further increased amyloid pathology in high AD stages (Fig. 3b). Within high neuropathological brain samples, the PrSr and PAS regions were revealed to have the highest levels of amyloid pixel density, followed by A36, MEC and A35 (Fig. 3b). Classifying different types of amyloid pathology in hippocampal regions revealed differences in pathology progression (Fig. 3c-e). APP expression appears to be higher in A36 and MEC in high cases (Fig. 3c). Diffuse amyloid is elevated in high cases in PrSr and PAS (Fig. 3d). Dense plaques also increased in most regions from low to high with A36 and A35 regions being the highest in high cases (Fig. 3e). Compared to dense amyloid, diffuse amyloid was significantly higher in the PAS and PrSr (Fig 3f3-4, PAS W = 665, p = 0.0021, PrSr W = 532, p = 0.0002). Although there was no significant difference in dense vs diffuse amyloid pathology in DG at the regional level, further analysis found laminar differences where the molecular layer of the DG (DGmo) had higher dense and diffuse plaque density compared to the polyform layer (DGpf) in high (Fig. 3g2 and 3g4; high diffuse DGmo vs DGpf W = 248, p = 0.0018; intermediate diffuse W = 226, p = 0.015; high dense W = 258, p = 0.0025).

**Fig. 3.**
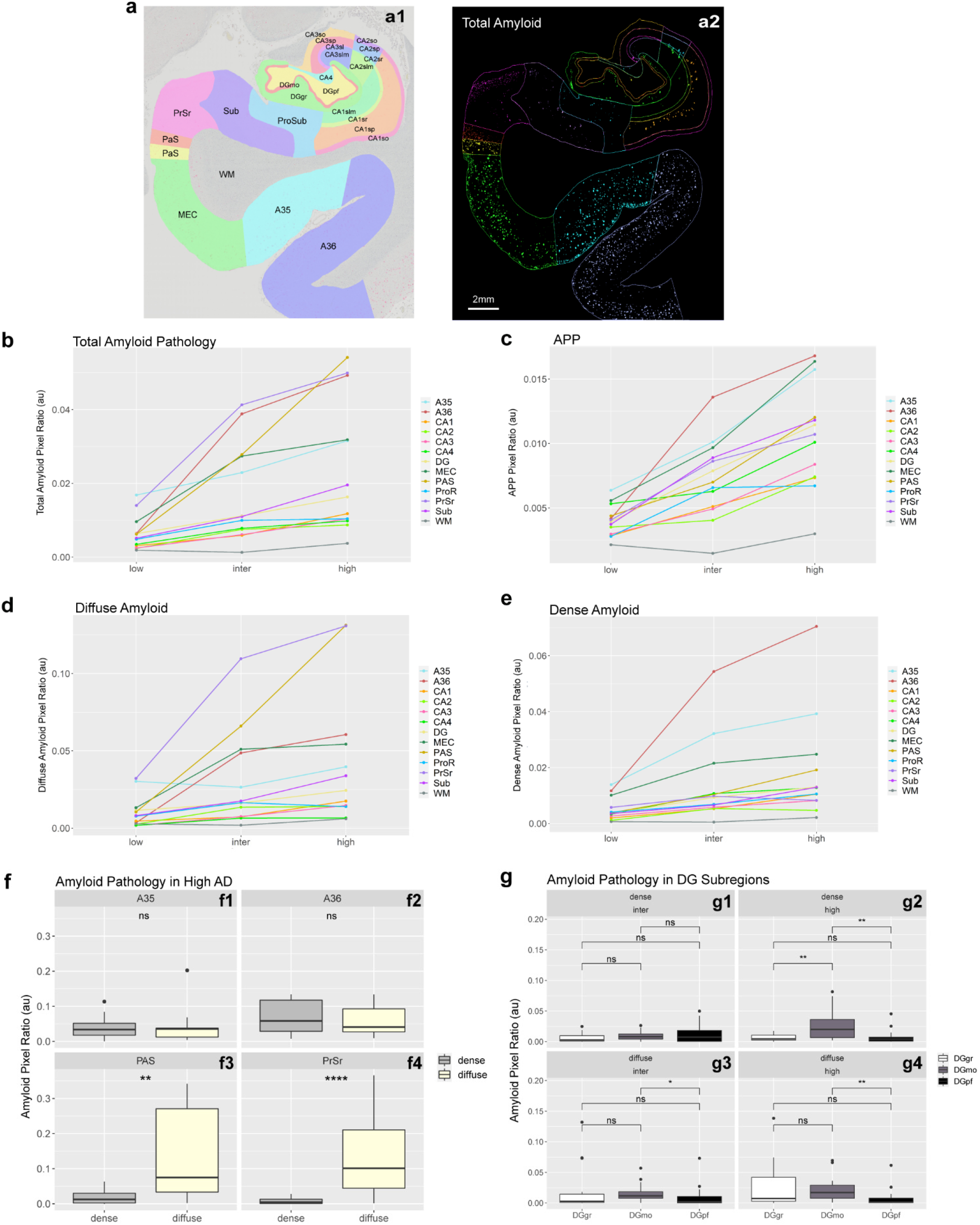
Subregional quantification of amyloid pathology. Manually segmented hippocampal subregions (a1) with corresponding ML extracted amyloid pixels (a2). Normalized individual subregional total amyloid data across low, intermediate and high cases (b). APP (c), diffuse (d), dense (e) pixel ratio in hippocampal regions in all stages. Dense and diffuse amyloid in A35 (f1), A36 (f2), PAS (f3) and PrSr (f4). Dense (g1-g2) and diffuse (g3-4) amyloid in DG subregions. Box plots show median (box center line), interquartile range (IQR, bounds of box), minimums and maximums within 1.5 times the IQR (whiskers), and outliers (points beyond the whiskers). p ≥ 0.05 was considered not significant (ns); *p < 0.05, **p < 0.01, ***p < 0.001.

We next assessed Gallyas positive NFTs in the same hippocampal regions (Fig. 4a). NFTs were separated from artifact staining and blood vessels (Fig. 4a1). When compared to low AD, high cases had significantly elevated levels of NFTs within whole hippocampal/medial temporal lobe sections (Fig 4b, W = 170, p = 0.003). At the subregional level, NFTs were primarily localized to the MEC, PrSr and PAS in low cases (consistent with early Braak stages; Fig. 4c). Within high AD patients A36 and CA4 contained the most NFTs, although Gallyas positive pixels were present within all anatomical subregions (Fig. 4c). NFTs were significantly higher in males in the A35 region in intermediate cases (Fig. 4d, W = 7, p = 0.042). We then assessed amyloid and Gallyas colocalization to identify extracellular neuritic plaques (Fig. 4e, n = 25-35). Neuritic plaques were significantly higher in intermediate cases compared to low (Fig. 4f, W = 8, p = 0.029). A36 and MEC were found to contain the highest density of neuritic plaques in high cases (Fig. 4g). In addition Gallyas positive NFTs were not correlated with amyloid pathology (Supplementay Fig 1a-b).

**Fig. 4.**
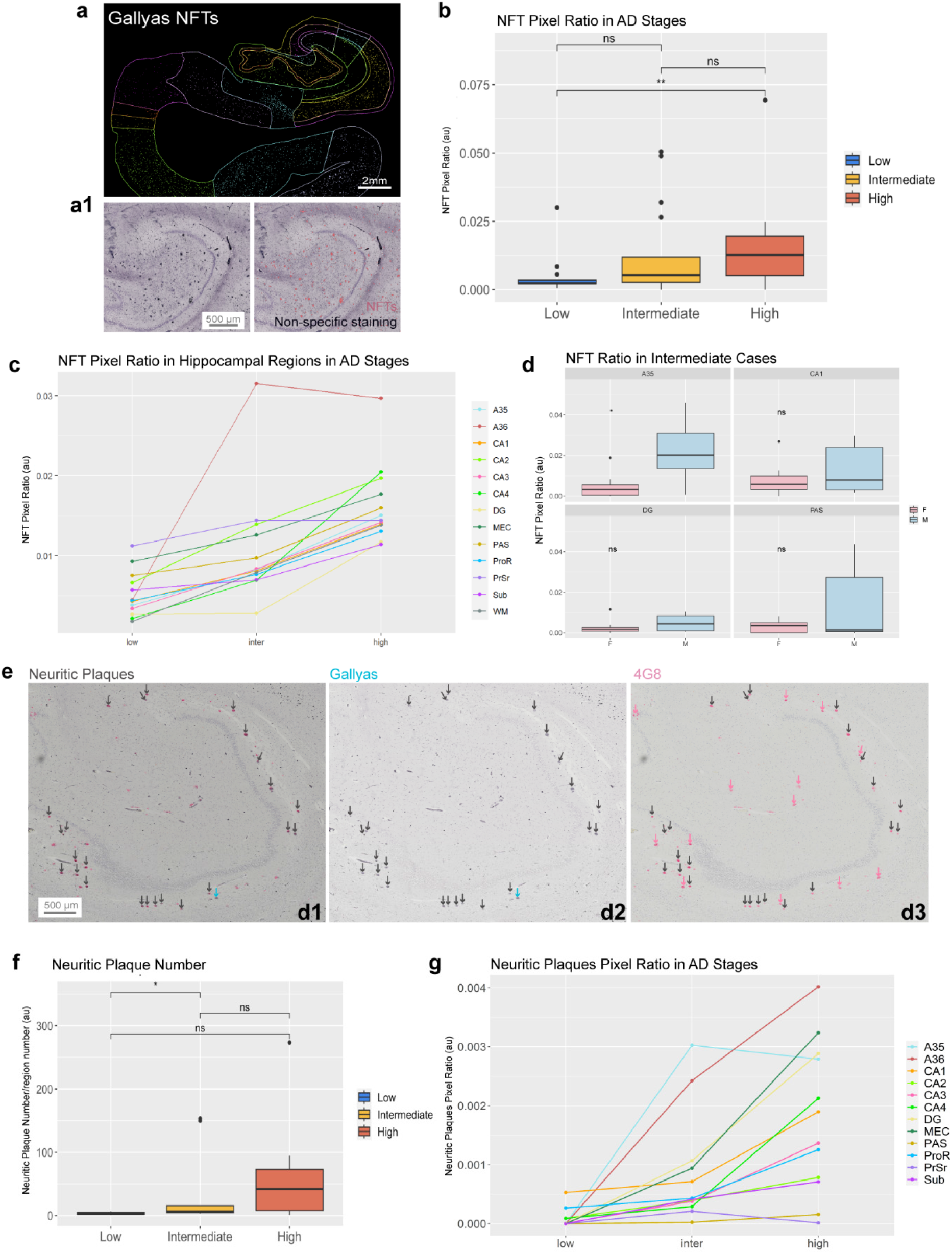
Gallyas positive NFTs in hippocampal and medial temporal lobe subregions (a). Identifying NFTs excluding non-specific staining (a1). NFTs across different AD stages (b). Regional differences in NFTs - A36 and CA4 contained the most NFTs although all anatomical subregions contained some NFT staining (c). Regional sex differences in NFTs (d). Example 4G8 amyloid and Gallyas NFT colocalized neuritic plaques (e). Neuritic plaques were significantly higher in high cases compared to intermediate cases (f). A36 and MEC were found to contain the highest density of neuritic plaques in high cases (g). Box plots show median (box center line), interquartile range (IQR, bounds of box), minimums and maximums within 1.5 times the IQR (whiskers), and outliers (points beyond the whiskers). p ≥ 0.05 was considered not significant (ns); *p < 0.05, **p < 0.01, ***p < 0.001.

To analyze the effect of ApoE allele genotype on AD pathology types, cases were further separated into ApoE4 carriers (4^+^) and ApoE4 non-carriers (4^-^). ApoE4 carriers had elevated levels of diffuse amyloid in high cases compared to non-ApoE4 carriers (Fig. 5a p = 0.038). Interestingly, we found that ApoE4 non-carriers 4^-^ had more dense plaques (Fig. 5c) and dense plaques were on average larger than the average dense plaque in ApoE4 carriers in high AD (Fig. 5d, W = 20, p = 0.041).

**Fig. 5.**
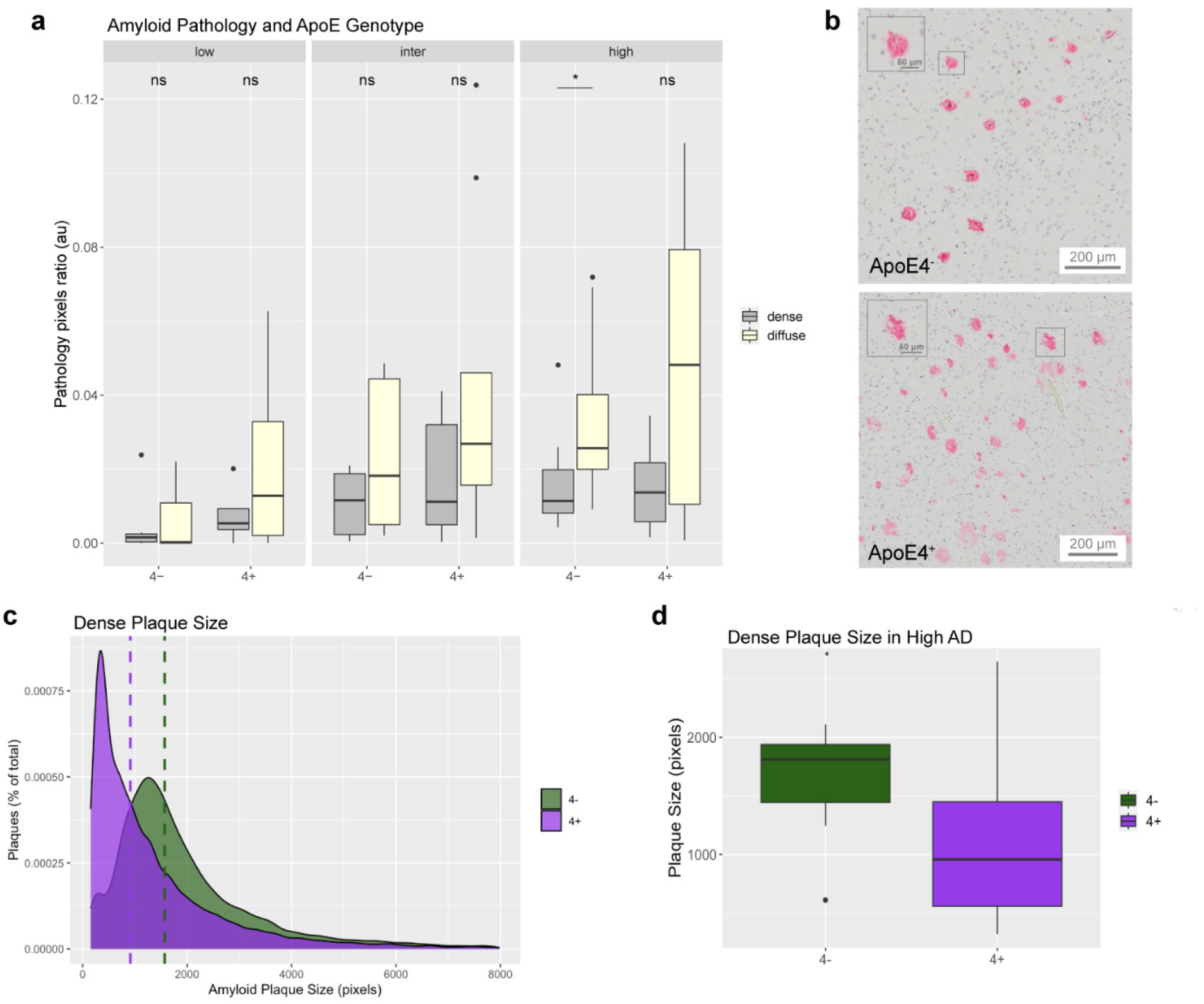
Overall diffuse and dense amyloid in ApoE4 carriers 4^+^ and non-carriers 4^-^ (a). Example images of dense plaques in ApoE4 non-carriers 4^-^ and carriers 4^+^ (b). Dense plaque size histogram across all stages (c). Dense plaque size is higher in non-carriers 4^-^ in high AD (d). Box plots show median (box center line), interquartile range (IQR, bounds of box), minimums and maximums within 1.5 times the IQR (whiskers), and outliers (points beyond the whiskers). p ≥ 0.05 was considered not significant (ns); *p < 0.05, **p < 0.01, ***p < 0.001.

To investigate the potential role of microglia in the expression of dense vs diffuse amyloid pathology and plaque size, we stained and analyzed a subset of adjacent hippocampal sections for Iba1^+^ microglia (Fig. 6a-b). We found sub-regional differences in microglia density that decreased from low to intermediate AD stages (except for MEC). Iba1^+^ microglial staining was highest in CA2, CA1 and DG, with some variance across stages (Fig 6c). This data was taken from a subset of cases (n = 9). Iba1^+^ microglia did not correlate with amyloid or NFTs, except for the DG, which was positively correlated with dense, diffuse and APP amyloid pathology (Fig. 6d, dense *R* = 0.9, p = 0.008; diffuse *R* = 0.8, p = 0.02; APP *R* = 0.9, p = 0.002). This is a preliminary assessment of microglia in the human hippocampus and further analysis is required to delineate more precise anatomical segmentation data for Iba1^+^ microglia.

**Fig. 6.**
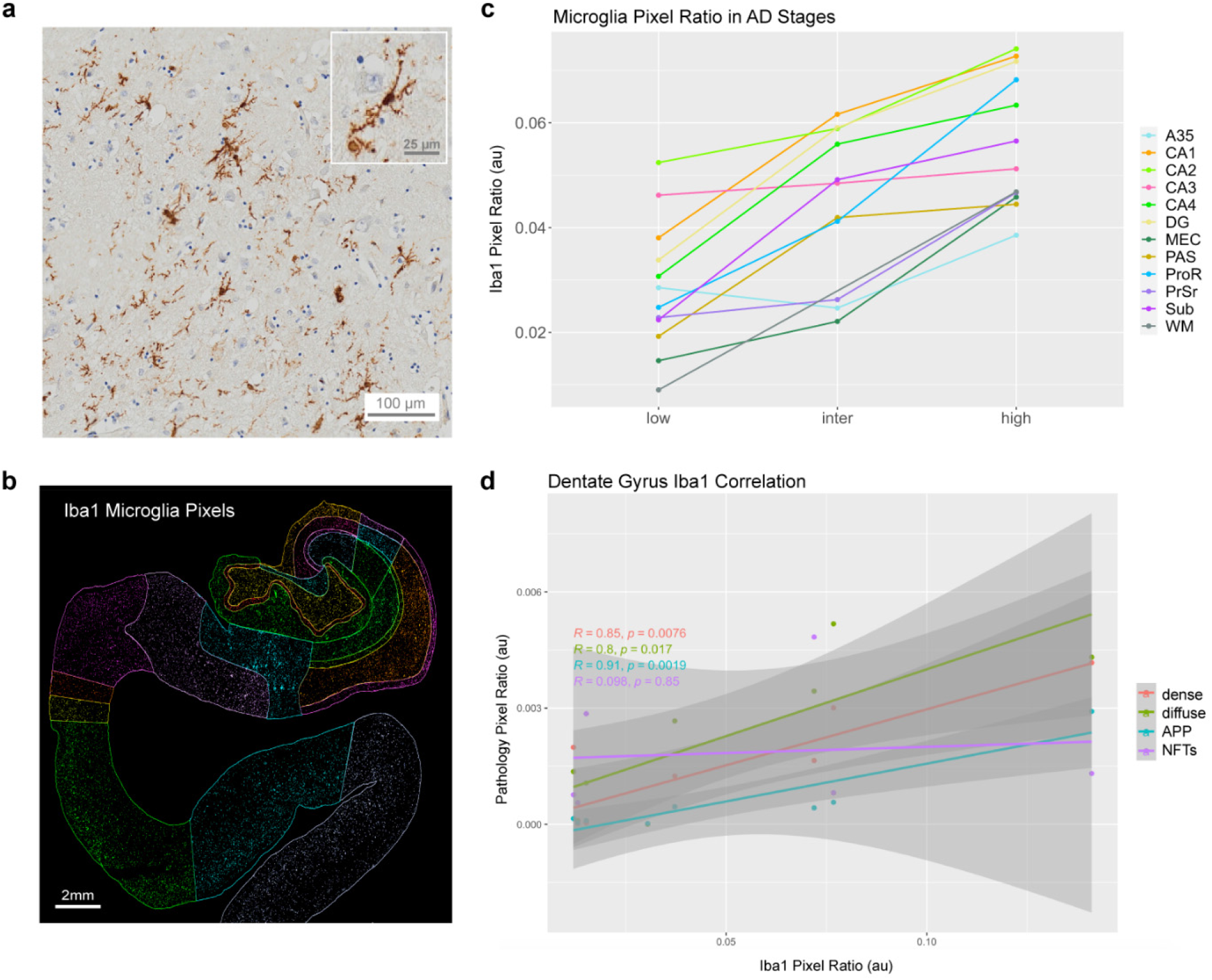
Iba1^+^ microglia staining example (a) and segmented regional staining in the hippocampus and medial temporal lobe (b) Iba1 staining was highest in the DG in low and intermediate cases (c) this data was taken from a subset of cases (n = 9). Iba1^+^ microglia did not correlate with amyloid or NFTs, except for the DG, which was positively correlated with dense, diffuse and APP amyloid pathology (grey inset shading shows IQR) (d).

Finally, we investigated whether AD pathology types in the medial temporal lobe were related to cognitive impairment, specifically the memory part of CDR. Gallyas positive NFTs and Iba1 microglia were not correlated with memory impairment, but dense and diffuse amyloid were positively correlated (Fig. 7a, dense *R* = 0.3, p = 0.03; diffuse *R* = 0.32, p = 0.02). In patients that had clinically diagnosed memory impairment, dense and diffuse amyloid was higher compared to mild or non-impaired patients (Fig. 7b, dense W = 318, p = 0.012; diffuse W = 335, p = 0.004). Regional specificity in combined stages: ProR, Sub, CA1, CA3, DG dense and diffuse were higher in impaired vs non-impaired individuals. Diffuse was also higher in impaired vs non-impaired cases in A36, MEC, PAS, PrSr. Dense was higher in A35 impaired individuals (Fig. 7c). In ApoE4 non-carriers both diffuse and dense amyloid were higher with memory impairment (Fig. 7d, dense W = 105, p = 0.025; diffuse W = 107, p = 0.02). Further, general linearized model plots showed that only diffuse pathology was correlated with CDR memory scores (Fig. 7e-g, p = 0.036).

**Fig. 7.**
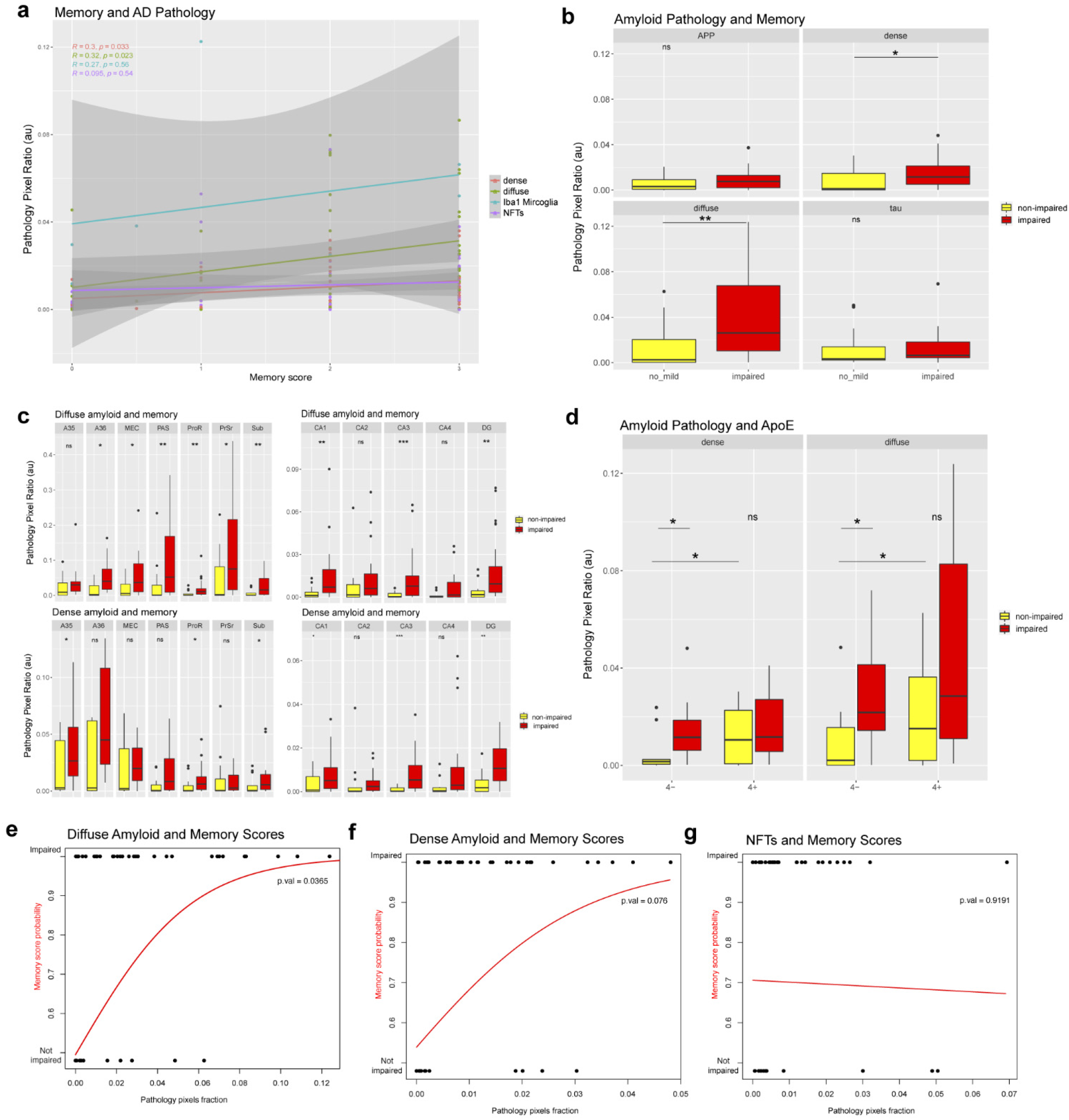
Pathology correlations with CDR memory scores (grey inset shading shows IQR) (a). APP, dense and diffuse pathology in mild/not impaired (yellow) and impaired (red) individuals (b). AD regions in mild/not impaired (yellow) and impaired (red) individuals across dense and diffuse amyloid (c). In ApoE4 non-carriers both diffuse and dense amyloid were higher with memory impairment (d). General linearized model (GLM) plots for diffuse (e), dense (f) and NFT (g) pathology. Box plots show median (box center line), interquartile range (IQR, bounds of box), minimums and maximums within 1.5 times the IQR (whiskers), and outliers (points beyond the whiskers). p ≥ 0.05 was considered not significant (ns); *p < 0.05, **p < 0.01, ***p < 0.001.

## Discussion

Accumulation of pathology within the hippocampus is a defining characteristic of AD progression and disease staging. Pathological assessment of AD brain tissue is based on qualitative scoring, which cannot fully characterize the distribution of pathology within anatomical brain subregions. Here, we demonstrate the development of a ML-based quantification and segmentation approach using openly available software that can rapidly quantify and classify the distribution of AD pathology within distinct subregions of the hippocampus and medial temporal lobe. Using this pipeline, we quantified the distribution of 4G8 amyloid, Gallyas-labeled NFTs, and co-registered neuritic plaques within the hippocampus and medial temporal lobe. Across 51 USC ADRC patients, significant heterogeneity was evident in different types of AD pathology and how they are expressed across medial temporal lobe regions as well as how they contribute to cognitive impairment. Among our findings, we identified that diffuse amyloid accumulation is the primary contributor to overall amyloid burden and was significantly correlated to CDR-assessed cognitive impairment.

4G8 antibody staining of intracellular APP, dense plaques, and diffuse amyloid demonstrates how complex labeling patterns can be identified and classified into distinct morphological structures by machine learning algorithms. PrSr and PAS regions were identified as areas with the highest levels of diffuse amyloid pathology, whereas dense plaques and APP were most concentrated in the A36 cortical region. Diffuse amyloid pathology in the PrSr has previously been described as having a ‘lake-like “ appearance^25^.

Interestingly, overall amyloid and NFTs were not correlated, but significant regional correlations did exist (supplementary figure 1a-b) highlighting the importance of not overlooking regional differences in AD. As expected, our data shows that NFTs increase across disease stage, but at different rates within anatomical brain regions. A36 and CA4 contained the most NFTs in high AD patients, but whereas A36 was also highly elevated in intermediate patients CA4 was relatively low at that stage. Another paper showed that entorhinal, CA1, CA3, and CA4 p-tau deposition levels are significantly correlated with Aβ burden, while CA2 p-tau was not^26^.

Microglia are primary inflammatory cells in the brain and changes in their localization or morphology are now known to be involved in disease initiation and progression^27^. Interestingly, we found that diffuse amyloid was higher and dense plaque size was reduced in ApoE4 carriers. Recent evidence suggests that microglia, specifically, are responsible for compacting diffuse amyloid into dense plaques^28^. 5xFAD mice which lack microglia develop diffuse and vascular amyloid pathology, but do not develop dense plaques like normal 5xFAD mice^29^. This suggests that compromised activities of microglia are involved in the reduction of dense plaque formation in our dataset. Microglia are neuroprotective, engulfing, and phagocytosing Aβ, but can also become activated, transitioning into disease-associated states, dependent on the presence of ApoE. Identifying exactly where and when they shift towards this damaging phenotype will undoubtedly aid therapeutic strategies for AD.

While we did not find any correlations between NFTs and Iba1, we did find significant correlations between amyloid pathology and Iba1 in the DG, but not other brain regions. Our results also show that the DG is a susceptible region for amyloid pathology, particularly diffuse amyloid. Specifically, the molecular layer subregion of the DG had higher levels of diffuse and dense amyloid pathology. The DG was also a site of high neuritic plaque burden. The high degree of colocalization between amyloid and tau NFTs suggests that DG neurons may be susceptible to neuritic plaque formation and consequently, neuronal damage^30^. It is important to further discern regional sex and ApoE differences in pathology to improve diagnostic capabilities. It would also be necessary to further investigate microglial distribution in high and non-AD cases, but our data does suggest that in subset of AD cases Iba1-positive microglial density is higher in the DG. Iba1-positive microglial density was reported to be higher in the mouse DG compared to other hippocampal regions^31^. And another study found that degeneration of microglia occurred in the DG of AD patients^32^. The DG might be a brain region with heightened microglia presence affected by AD. Whether or not this effect is beneficial or detrimental remains to be determined. The DG has a significant contribution to the formation of episodic memory^33–35^, as well as being a site of neurogenesis throughout the human lifespan^36^. Alongside our data, this implicates the accumulation of possibly impaired microglia in the DG, which might affect adult neurogenesis in the early stages of AD. It should be noted that the development of AD could be due to either excessive or deficient microglial activation.

Aligning our data with clinical data revealed that dense and diffuse amyloid were significantly elevated in patients that had diagnosed memory impairment, but NFT density was not. Walker et al., show that cognitive impairment was correlated with overall p-tau burden. Clinical data in the form of CDR Global Scores correlated with p-tau and not amyloid burden^26^. Our data did not show any correlations with Gallays labeled NFTs, but it did reveal that regional differences are present between impaired and non-impaired individuals, suggesting a link between pathology and cognition that might be detected before the disease becomes too advanced. Additionally, a recent study has shown that NFT maturity may play an important role as ghost tangles, but not mature tangles or pre-tangle tau, were correlated to cognitive deficits^37^.

In contrast to our findings of a significant correlation of dense and diffuse amyloid with cognitive impairment, a recent study found that dense Aβ plaques, but not diffuse plaques, in the hippocampus were linked to worse AD neuropathology. Multiple differences between our studies could explain these outcomes including differences in amyloid antibody staining (4G8 vs Amyloid ꞵ), cognitive testing (CDR vs ECoG), patient demographics (primarily white vs. Asian), and quantification approach (machine learning vs manual counts). A potential explanation could be that the previous study analyzed dense plaques whereas our study focused on diffuse amyloid which can often aggregate as a ‘plaque ‘ but also larger aggregates like in the PrSr^38^.

## Conclusions

Overall, we report quantitative neuropathological assessment of 51 AD patients with novel anatomical region-specific differences as well as sex, ApoE genotype, and microglial heterogeneity. Understanding where and when AD pathology is present in different anatomical brain regions and which cell types are susceptible to neurodegeneration is critical to developing effective treatment and prevention strategies. The hippocampus, amygdala, and entorhinal cortex are anatomical areas associated with the pathology of AD, although these medial temporal structures are not where neuropathology correlates best with dementia at any stage of the disease. Other areas across the brain contain pathological aggregates and play an important role in disease progression. Some studies have reported that as many as 25% of AD cases appear to spare the hippocampus^8,39^ so it will be important to continue classifying other brain regions such as the frontal cortex. With more high-throughput digital pathology and deep learning analysis, computational pathology has the potential to improve diagnostic quality, clinical workflow efficiency, with a view to create personalized treatment plans for patients. Going forward it will be important to align this ML-based segmentation approach with patient demographic information, clinical records, and ultimately functional MRI imaging, to create a longitudinal perspective of disease progression. We believe this approach can be utilized to improve pathological diagnosis, classification, prediction, and prognosis of AD.

## Abbreviations

AD: 
ADRC: 
NFTs: 
ApoE: 
ADRC: 
ROI: 
QIT: 
DG: 
MEC: 
PAS: 
ProR: 
PrSr: 
Sub: 
APP: 
ML: 
CDR: 

## Acknowledgements

This work was supported by NIH grants: K01AG066847 (author MSB), P30-AG066530 (author DH), as well as the Epstein Family Foundation Alzheimer’s Research Collaboration Gene Therapy Program. We would like to acknowledge the USC ADRC clinicians, researchers, patients, and administrators whose efforts have made this study possible. Slides for this study were obtained from the Alzheimer’s Disease Research Center Neuropathology Core, NIA AG066530. RPC’s contributions were supported by grant number 2020-225670 from the Chan Zuckerberg Initiative DAF, an advised fund of Silicon Valley Community Foundation. MP’s contributions were funded by an NIA supplementary grant 3P30AG066530-02S1.

---

**Supplementary Figure 1.**
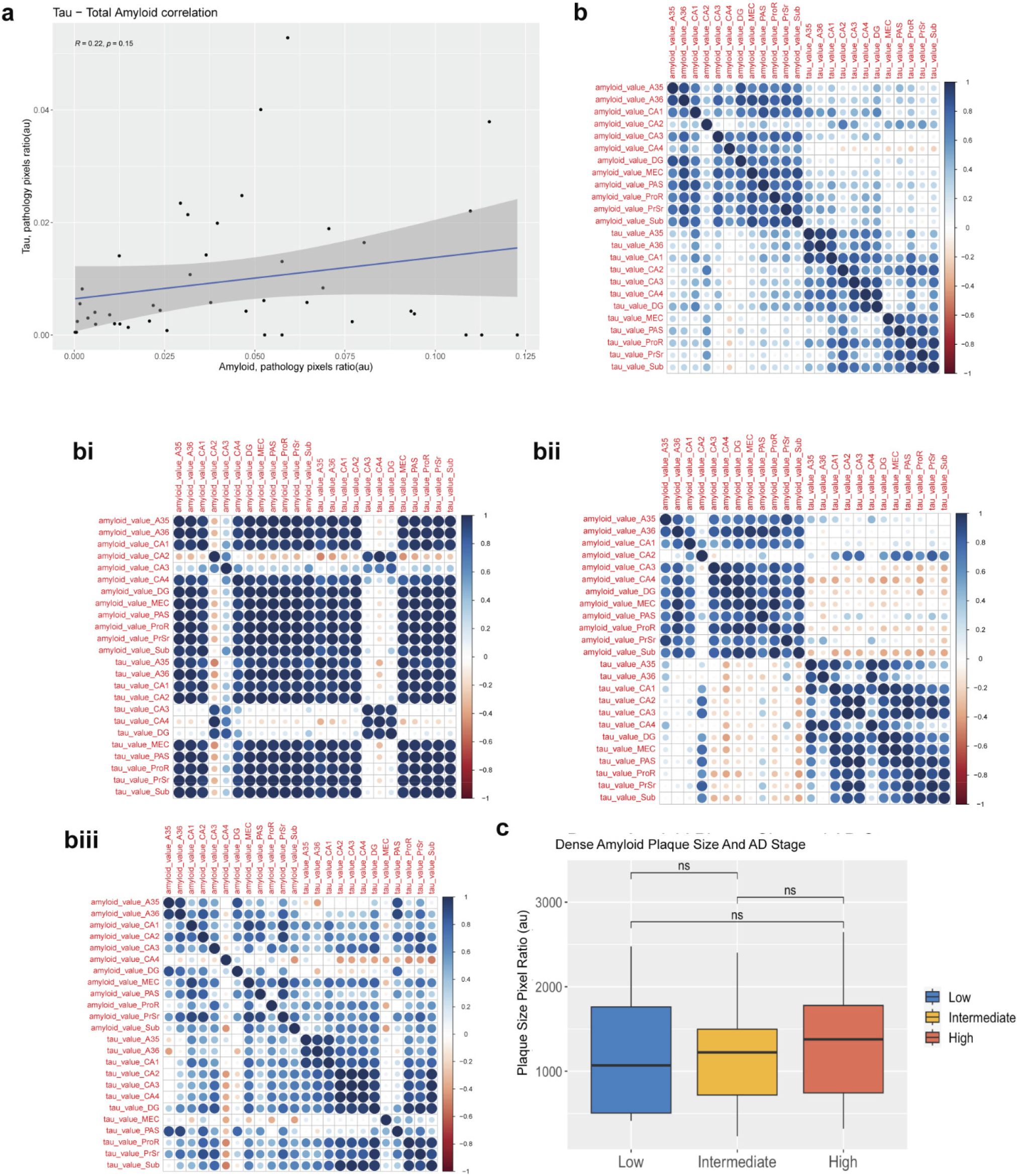
Linear regression of total amyloid and Gallyas NFT correlation plot (a). Regional differences in total amyloid and Gallyas NFTs. A correlation matrix was created to examine the relationship between amyloid and tau pathology in hippocampal subfields. The amyloid and Gallyas NFT load for each case were used to calculate Pearson’s correlation coefficients. The correlation matrix was then created using RStudio with circle size representing the magnitude of the correlation coefficient and color indicating the direction of the correlation (red for negative correlation and blue for positive correlation) (b). Low (bi), intermediate (bii), and high correlation matrices (biii). Dense amyloid plaque size in AD stages (c). Box plots show median (box center line), interquartile range (IQR, bounds of box), minimums and maximums within 1.5 times the IQR (whiskers), and outliers (points beyond the whiskers). p ≥ 0.05 was considered not significant (ns); *p < 0.05, **p < 0.01, ***p < 0.001.

